# Minimal overlaps in responses to insecticides between pollinator species

**DOI:** 10.1101/2025.05.19.654787

**Authors:** Alicja Witwicka, Federico López-Osorio, Courtney May, Yeahji Jeong, Yannick Wurm

## Abstract

Insecticides are important for protecting crops from agricultural pests, yet their use inadvertently drives global declines in beneficial insects, including pollinators. Insecticide regulation relies on the assumption that model bee species adequately represent responses to exposure across the diversity of insect taxa, despite over 479 million years of evolutionary divergence and a limited understanding of how exposure effects vary across insect orders. Here, using comparative whole-brain transcriptomics, we show differences in molecular responses to modern insecticides across evolutionary distant pollinator lineages: butterflies, flies, and bees. Within each of the four species studied, different insecticides (sulfoxaflor and clothianidin) triggered broadly similar gene regulatory responses. In surprising contrast, exposure impacts differed sharply among species, with no genes or pathways being consistently affected. Strikingly, we found that sulfoxaflor, approved based on its supposed safety for bees, causes more disruptions in non-bee pollinators than the restricted neonicotinoid clothianidin, revealing a critical blind spot in current assessment protocols. Our findings demonstrate that over large evolutionary timescales, species-level differences can outweigh variations in insecticide chemical structures in shaping the effects of insecticide exposure. These interspecific differences likely reflect distinct physiological and metabolic traits shaped over tens to hundreds of millions of years. Together, our findings highlight the urgent need to reevaluate insecticide safety assessments to incorporate phylogenetic diversity, potentially explaining why regulatory efforts have failed to halt pollinator declines despite stricter testing requirements.

**Significance Statement:** Insecticide regulations are based on a narrow set of test species, overlooking the evolutionary and functional diversity among pollinators. Here, we show that molecular responses to insecticides differ more between species than between chemicals, even when targeting the same neural receptors. Strikingly, an insecticide considered safe for bees caused greater molecular disruption in non-bee pollinators. Our findings challenge current assumptions in pesticide risk assessment and suggest that using surrogate species can misrepresent broader ecological risks. By revealing species-specific vulnerabilities rooted in millions of years of divergent evolution, this study underscores the need for a more inclusive and phylogenetically informed approach to protecting pollinator biodiversity.

## Introduction

Modern intensive agriculture typically involves heavy use of insecticides (Schulz et al., 2021; Tang et al., 2021) which have detrimental effects on beneficial insect pollinators (Butler, 2018; Goulson, 2013; Nicholson et al., 2024; Rundlöf et al., 2015; Tsvetkov et al., 2017; Woodcock et al., 2017). Due to these harmful impacts, certain previously authorised insecticides are locally restricted or banned, leading to the introduction of alternative compounds. However, these replacements often prove equally damaging (Siviter & Muth, 2020; Siviter et al., 2018, 2024). This cycle persists because despite similar biological modes of action, insecticides fall into distinct groups based on their chemical structures (Casida, 2018). This classification often leads to regulatory decisions for each group, for example, the restriction on neonicotinoids in the EU (Siviter et al, 2023). Furthermore, insecticide safety assessments rely primarily on the honey bee (*Apis mellifera*) as a model surrogate species for the enormous diversity of pollinating insect groups (Franklin & Raine, 2019). The inadequacy of this surrogacy strategy becomes evident as approved insecticides continue to adversely impact wild pollinator populations (Guzman et al., 2024; Nicholson et al., 2024; Wagner et al., 2021; Woodcock et al., 2017), highlighting a critical need to understand how insecticide responses vary across insect species.

With a history spanning over 479 million years (Misof et al., 2014; Peña-Kairath et al., 2023), insects have evolved diverse adaptations for metabolising foreign chemicals of natural or synthetic origin, suggesting that species might respond differently to insecticides (Nauen et al., 2022; Raine & Rundlöf, 2024). However, most modern insecticides target highly conserved nicotinic acetylcholine receptors (nAChRs) and the acetylcholine signalling system in all insect brains (Casida, 2018), raising the possibility of uniform cascading responses across species. Therefore, two competing scenarios are possible: (I) detoxification gene families shaped by distinct selective pressures underlie species-specific molecular responses to insecticides, or (II) the conserved nature of acetylcholine signalling and its downstream pathways results in broadly similar molecular responses across species. We investigated these hypotheses by comparing brain gene expression responses to insecticide exposure across four diverse pollinating insect species.

We performed whole-brain transcriptomic analyses to dissect the effects of two prevalent insecticides (clothianidin and sulfoxaflor) on four pollinating insect species representing three orders: a widely spread butterfly species, the painted lady (*Vanessa cardui*), the agriculturally significant green bottle fly (*Lucilia sericata*), the solitary red mason bee (*Osmia bicornis*), and the social buff-tailed bumble bee (*Bombus terrestris*). These species span distinct evolutionary lineages, life histories, and ecological roles, providing insight into broader patterns of species-specific responses to insecticides. We exposed all insects to 4.4 ppb of both insecticides, a concentration commonly found as residuals in pollen and nectar (Wood & Goulson, 2017). Butterflies and bees were exposed over 12 days, while flies, because of their shorter lifespan, were exposed over 8 days (see Materials & Methods).

Our findings reveal substantial variation in how different pollinator species respond to insecticide exposure, underscoring the complexity of assessing insecticide risks. The magnitude of gene expression changes varied markedly across species, affecting the number of differentially expressed genes and the direction of regulation. We found no consistent responses at the gene or pathway level across species. Despite these differences, gene expression responses to different insecticides within each species were highly consistent, with over 70% of differentially expressed genes showing similar regulation across treatments. These results suggest that while insecticides act on conserved neural pathways, the downstream molecular responses are highly species-specific, challenging the validity of single-species surrogacy in risk assessments.

## Results & Discussion

### Different insecticides affect overlapping gene sets within-species

We detected contrasting effects of sulfoxaflor and clothianidin across species. Clothianidin triggered differential expression in 2,248 genes in the bumble bee and 759 genes in the solitary bee, while sulfoxaflor caused minimal changes in both bee species. By contrast, sulfoxaflor induced differential expression in 12 times more genes in butterflies and in eight times more genes in flies compared to clothianidin (Figure 1, Supplementary Tables 1 and 2).

**Figure 1.**
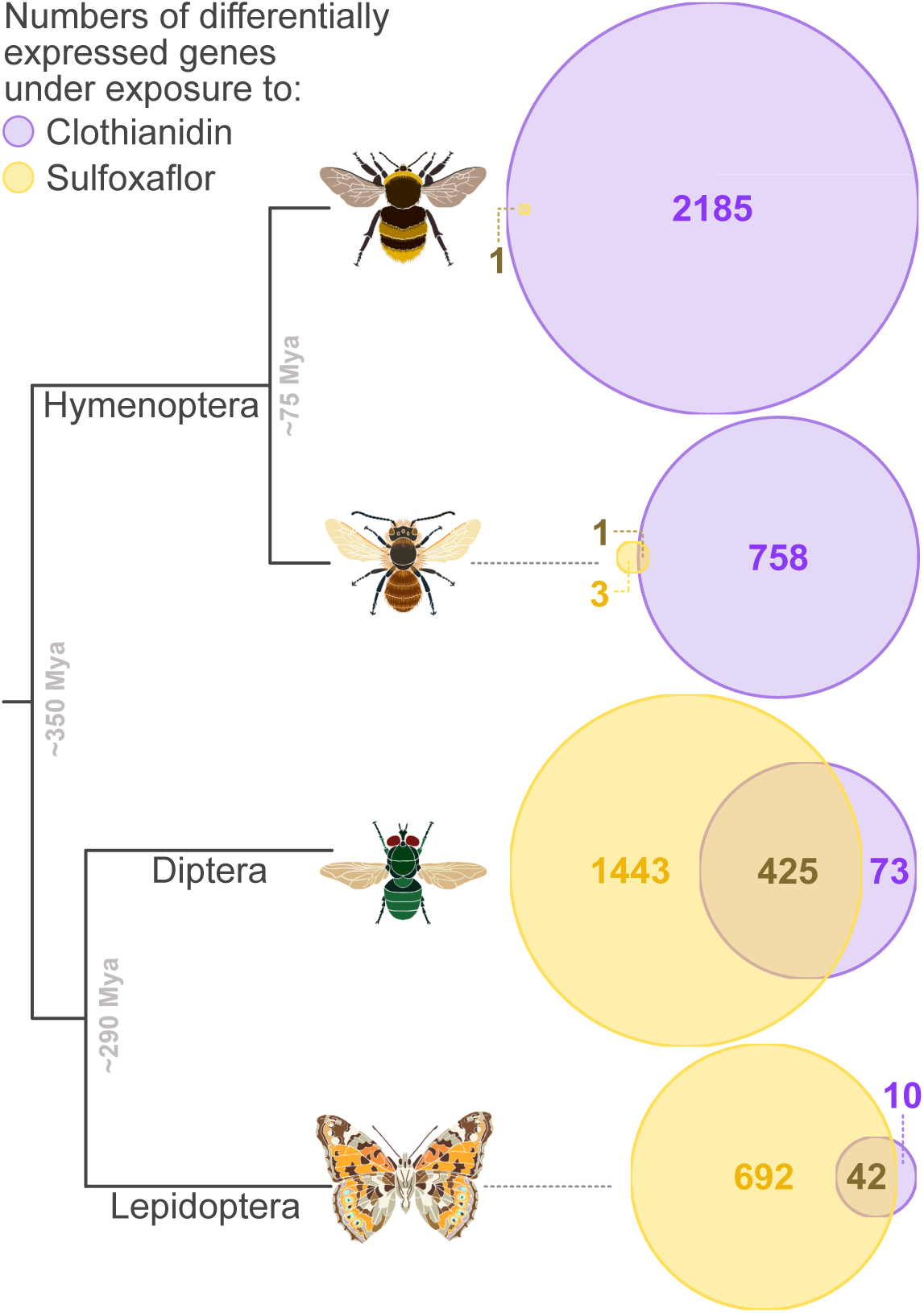
Euler diagrams representing the number of differentially expressed genes under clothianidin (purple) and sulfoxaflor (yellow) exposure in the studied species and the species’ topology and estimated divergence times.

Sulfoxaflor has been implicated in reproductive effects in bumble bees (Siviter et al., 2018), leading to its restriction by the European Commission. However, subsequent studies have yielded inconsistent findings on its impacts on bees (Schwarz et al., 2022; Siviter et al., 2019; Straw et al., 2023), while data on non-bee species remain scarce. In other insects, sulfoxaflor has been linked to reduced fertility and oviposition in *Harmonia axyridis* (Dai et al., 2021), mitochondrial dysfunction in *Chironomus kiinensis* (Liu et al., 2021), and toxicity comparable to imidacloprid in *Locusta migratoria* (Parkinson et al., 2020). Our data show that at molecular level, sulfoxaflor causes more pronounced effects than a commonly used neonicotinoid in at least some non-bee species.

We observed overlaps of differentially expressed genes under exposure to clothianidin and sulfoxaflor within species. In the fly, 85%, and in the butterfly, 81% of genes differentially expressed under clothianidin exposure were also affected by sulfoxaflor (hypergeometric test P-values < 10^−11^ for both). The direction and amplitude of expression changes in these genes were highly correlated between treatments in both species (Figure 2A and B), indicating that the two insecticides cause similar molecular disruptions in these insects. To further investigate this pattern, we identified gene modules with coordinated expression shifts under exposure to both treatments. Because genes often exhibit coordinated expression patterns, even when individual gene changes do not reach statistical significance, we hypothesised that overlapping gene modules would be affected by both treatments. In both species, two gene modules showed correlated responses to both insecticides (Supplementary Tables 3 and 4). Notably, 68% of genes in flies and 33% in butterflies significantly differentially expressed only under sulfoxaflor were part of these modules, exhibiting correlated shifts under both insecticides. Under clothianidin, these genes showed weaker expression changes that did not reach statistical significance (Figure 2C, D).

**Figure 2.**
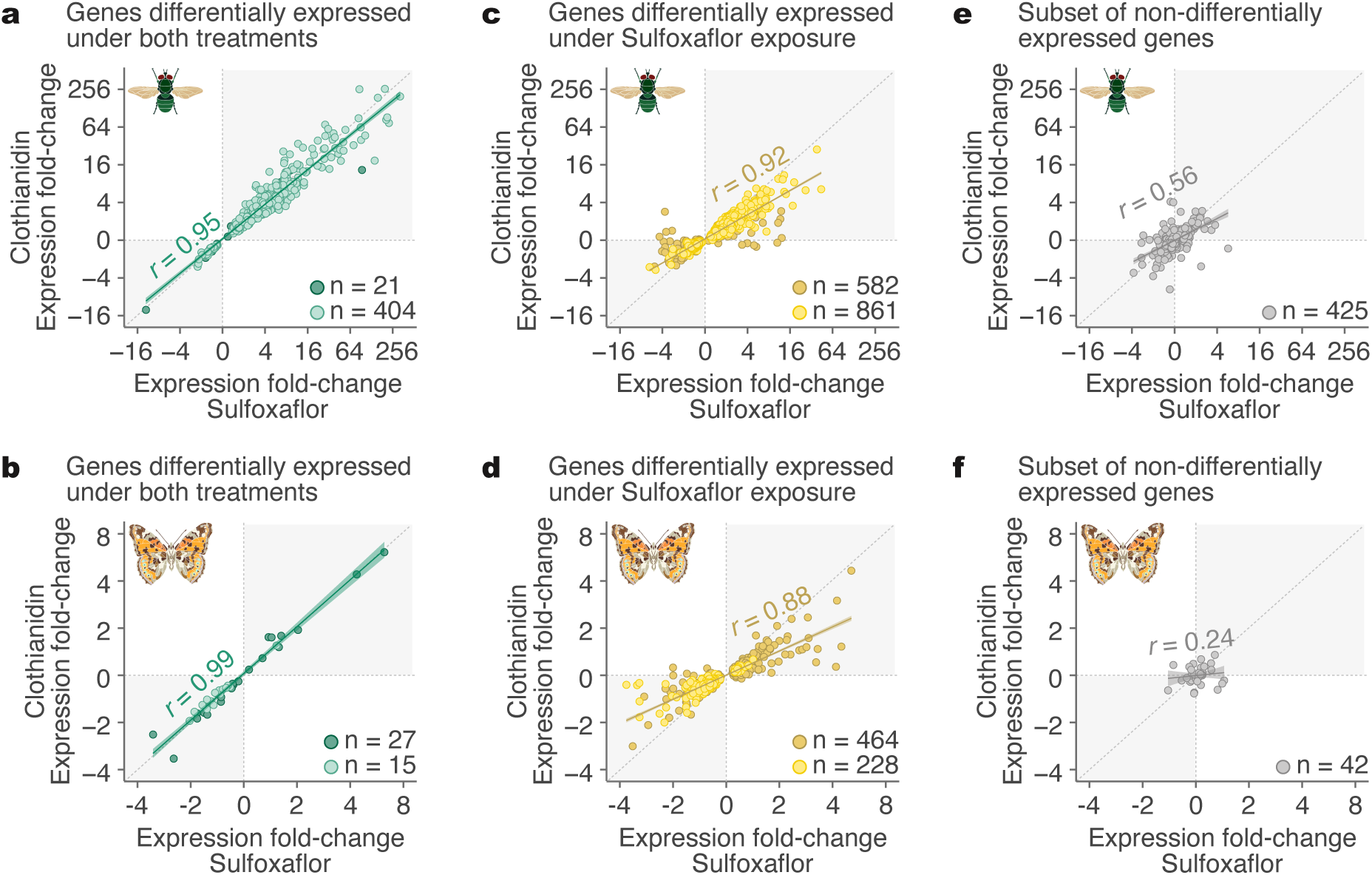
Regression plots indicating the similarities in the magnitude of expression fold-change in genes affected by the two insecticides. Genes differentially expressed under both clothianidin and sulfoxaflor in **A.** green bottle fly, and **B.** painted lady butterfly. Genes differentially expressed under exposure to sulfoxaflor but not clothianidin in **C.** green bottle fly, and **D.** painted lady butterfly. Despite the genes not being detected as differentially expressed, majority of them still shifts their expression in the same direction under exposure to both insecticides. The lighter coloured data points (**A-D**) indicate genes found in the gene modules identified as correlated with exposure to both insecticides. Random subsets of non-differentially expressed genes in **E.** green bottle fly and **F.** painted lady butterfly as a comparison. The number of genes in these subsets is equal to the number of the differentially expressed gene overlap for both species (in **A** and **B**). The Pearson correlation coefficient (*r*) indicates the strength of the linear relationship between the two treatments for each comparison.

Because sulfoxaflor caused minimal transcriptomic changes in bees, we could not robustly compare its effects with clothianidin. To test whether different insecticides cause overlapping responses in red mason bees, we exposed this species to acetamiprid, another neonicotinoid insecticide. We detected 27 differentially expressed genes, 71.5% of which overlapped with those affected by clothianidin exposure, exhibiting correlated direction and amplitude of expression changes across both treatments (hypergeometric test P-value < 10^-22^, Supplementary Figure 1). We previously showed overlapping effects of acetamiprid and clothianidin in bumble bees (Witwicka et al., 2025). We further compared our results to a study on imidacloprid exposure (Bebane et al., 2019) to expand the number of compounds tested. We detected a 36.5% overlap in differentially expressed genes with those affected by clothianidin (hypergeometric test, P-value < 10⁻¹²). This overlap persisted despite variations in exposure duration and concentration that determine the molecular effects of exposure more than the compound type itself (Witwicka et al., 2025; Supplementary Figure 1). The altered expression of shared gene sets within individuals of one species exposed to different insecticides suggests similar biological mechanisms of action and downstream effects, albeit with varying intensities. These findings demonstrate that within species, certain gene networks are consistently affected by insecticide exposure, even when the responses vary in magnitude. This consistency supports our hypothesis that the conserved nature of cholinergic signalling plays a key role in determining exposure outcomes within species. While different insecticides may bind to receptors with varying affinities due to differences in subunit specificity ( Lu et al., 2022), the resulting downstream effects have broad similarity, differing primarily in intensity.

### Effects of insecticides differ between species

In each species, 23% to 29% of genes affected by the more impactful treatment (clothianidin for bees, sulfoxaflor for butterflies and flies) were amongst the conserved one-to-one orthologs shared across all four species (2,956 genes), suggesting that some conserved pathways are affected by the exposure. However, pairwise comparisons between species revealed minimal and non-significant overlaps amongst these genes (Supplementary Table 7). We found only one statistically significant overlap between the two bee species (hypergeometric test, p < 10^−5^; Figure 3A and B), but most of these genes also exhibited opposing directions of expression change (Figure 3C and D). Only one gene, *tafazzin*, a putative acyl transferase associated with altered metabolism of the mitochondrial phospholipid cardiolipin (Xu et al., 2006), was affected in all four species. However, it was downregulated in flies and butterflies under sulfoxaflor exposure but upregulated in both bee species under clothianidin exposure, furthering showing the differences in responses between species (Figure 3D and D). In all species, the proportion of enriched KEGG pathways unique to a single species ranged from 25% in the red mason bee to 62% in the painted lady butterfly. We detected no KEGG pathways enriched across all four species (Figure 3E). For example, oxidative phosphorylation was disrupted in both bumblebees and flies but remained entirely unchanged in butterflies. Likewise, immunity pathways were affected in both bees and flies, but the nature of these changes differed: bumble bees showed down-regulation of antimicrobial peptides, while flies showed up-regulation (Figure 3F).

**Figure 3.**
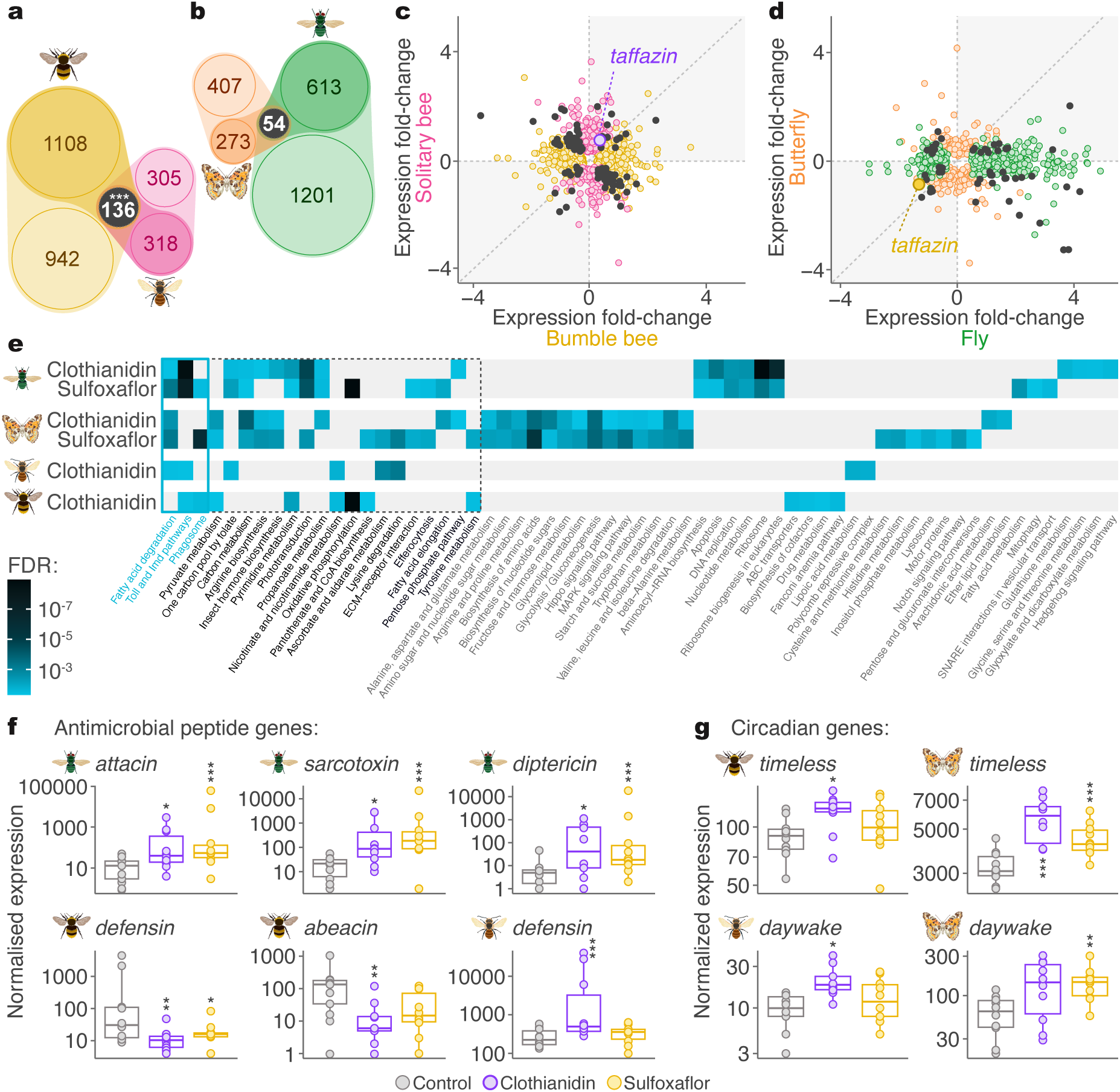
Between species changes in gene expression under insecticide exposure. **A.** Overlaps between differentially expressed one-to-one orthologs red mason bees and bumble bees, and **B.** in flies and butterflies. Lighter circles represent non-conserved differentially expressed genes. Darker circles indicate conserved one-to-one orthologs that were differentially expressed in only one species. Grey circles denote one-to-one orthologs that were differentially expressed in both species. **C.** Correlation of the magnitude of expression fold-change in one-to-one orthologs in bumble bees and red mason bees exposed to clothianidin, and **D. in** flies and butterflies exposed to sulfoxaflor. The grey data points indicate the overlapping affected one-to-one orthologs. The direction of expression change did not correlate between species. The blue data point represents the sole gene affected in all four species, though the direction of expression change varied across species. Outlier genes have been removed. **E.** Heatmap representing the KEGG pathways enriched under exposure to insecticides in all four species. Only three pathways were enriched in three species. No pathway was enriched across all four species. **F.** Changes of expression in antimicrobial peptides in flies, bumble bees and red mason bees. While the antimicrobial peptides were consistently upregulated in flies and red mason bees, they were downregulated in bumble bees. No changes were detected in butterflies. **G.** Changes of expression in circadian clock genes. *Timeless* gene was upregulated in butterflies and bumble bees. *Daywake* was upregulated in red mason bees and butterflies. No changes were detected in flies. Independence codes: * < 0.05; ** < 0.01; *** < 0.001.

Additionally, we detected changes in genes associated with circadian rhythm regulation in both bee species and the butterflies but not the flies (Figure 3G).

While similar detoxification mechanisms are employed to initially break down the toxic compound, the species-specific expansions and gene loss of detoxification gene families driven by differences in occupied habitats and subsequent evolutionary pressures likely determines the inherited capacity to cope with the exposure (Bass et al., 2024; Dermauw et al., 2020; Haas et al., 2022; Hayward et al., 2024; Manjon et al., 2018). At the same time, brain tissue shares fundamental molecular characteristics across species (Brawand et al., 2011; Cardoso-Moreira et al., 2019; Kannan et al., 2019; Supplementary Figure 2). Activation or blockage of nAChRs by insecticides triggers Ca²⁺ influx, disrupting neuronal signalling. This disruption likely leads to a limited set of biological alterations, explaining the overlapping pesticide effects within species. However, because of the interspecific differences in the composition of nAChRs, compounds may activate the nAChRs at different rates, thus determining the overall response dissimilarities across species (Jones et al., 2007; Witwicka et al., 2023). Therefore, while there likely is a limited number of effects caused by the nAChR activation in the brains, such impacts may be expressed at various strengths in various species depending on their ecological adaptations and base-level metabolic differences. Additionally, variations in anatomical and physiological traits among species, such as differences in neuron composition, body size, cuticle structure, lifespan, and feeding strategies, may amplify disparities in their reactions to exposure (Siddiqui et al., 2022).

### Rethinking insecticide safety assessment

Current methodologies for insecticide safety assessment focus primarily on the mortality rates, often neglecting sub-lethal effects that occur at field-realistic concentrations (Franklin & Raine, 2019; Schäfer et al., 2019; Topping et al., 2020). Our data show that exposure to insecticides can cause significant transcriptomic changes across species, even when survival rates remain relatively high (Supplementary Figure 2). These molecular changes likely underlie phenotypic and behavioural alterations (Crall et al., 2018; Goulson, 2013b; Siviter et al., 2021). The consistent yet species-specific responses to multiple insecticides reinforce the conclusion that evolutionary history, rather than chemical structure, shapes molecular responses. Although this study focuses predominantly on four species and two insecticides, the findings highlight consistent patterns of divergence in molecular responses, challenging the reliance on single-species models.

While changes in gene expression may reflect adaptive responses to environmental stressors, we would anticipate these responses to be more prominent under milder treatments, where metabolic systems are not yet overwhelmed and can activate protective pathways. However, insecticides with lower mortality rates largely elicit weaker effects on the same genes affected by the more lethal insecticide, suggesting a response driven mostly by toxicity rather than an adaptive mechanism (Supplementary Figure 3). Moreover, the brains of bumble bees and solitary bees, as well as other tissues in bumble bees (Malpighian tubules and hind femurs; Witwicka, 2024), appeared to tolerate sulfoxaflor exposure without exhibiting transcriptomic changes indicative of attempts to mitigate or counteract the exposure. These patterns suggest that the extent of transcriptomic changes, reflected in the number and intensity of differentially expressed genes, indicate insecticide toxicity levels and allow comparisons across compounds within a species. Thus, our results represent a compelling case for incorporation of comparative transcriptomics in insecticide safety assessments.

However, we reveal pronounced variability in molecular responses to insecticides even between relatively closely related bee species. One question remains, whether the variability in responses is correlated with phylogeny or ecological roles within ecosystems. With the rapid emergence of reference genomes for non-model species (Supple & Shapiro, 2018; Theissinger et al., 2023), comparative genomics offers a powerful opportunity to quantify and better understand this variation across taxa. Nonetheless, our findings suggest that regulatory frameworks based on model bee species alone are unlikely to reliably capture this variability in effects, given the evolutionary diversity of insects. Insecticide applications, regardless of refinement, will always pose challenges to biodiversity (Sánchez-Bayo, 2021; Wan et al., 2025). To address the diversity of processes affected across species, agricultural systems must transition from heavy reliance on insecticide applications to integrated practices that work with nature, reducing dependency on chemical controls and prioritising biodiversity as a cornerstone of sustainable food production.

## Supporting information

Sample Summary

## Acknowledgments

We were funded by the Natural Environment Research Council (NE/L00626X/1 and NE/S007229/1), the European Commission (H2020-MSCA-IF-2018-840185), and the Biotechnology and Biological Sciences Research Council (BB/T015683/1). This research used the Apocrita HPC facility, supported by QMUL Research-IT (http://doi.org/10.5281/zenodo.438045). We thank Professor Dave Goulson for his constructive input and thoughtful comments on the manuscript.

## Materials and Methods

### Experimental design

Females of bumble bees, solitary bees and painted lady butterflies were treated for 14 days. For the first two days of adjustment period, all individuals were fed control treatments. Subsequently, for the remaining 12 days, individuals were fed 4.4 ppb insecticide solution. Flies, because of shorter adult lifespan and considerably higher mortality upon exposure (> 95% mortality after exposure over 12 days to both sulfoxaflor and clothianidin) were subject to one day of adjustment period followed by 8 days of insecticide exposure to collect a sufficiently large sample size and measure sub-lethal effects of exposure on the transcriptome. Exposure solutions were provided *ad libitum* and changed every other day.

All insect species were subject to 8h of darkness and 16h of light. Insects were handled under red light to minimise flight activity during feeding. All insects were kept at 24℃ (± 1℃). All feeding happened between 11:00 am and 1:00 pm to minimise the effects of circadian rhythms on gene activity. All sampling took place between 1:00 pm and 2:00 pm. Upon the end of exposure, insects were placed in 2 ml screwcap cryo-tubes, rapidly submerged in liquid nitrogen, and stored at -80℃ for further processing. We obtained 10 replicates for each species and treatment group combination (30 replicates per species, 120 replicates total).

### Osmia bicornis

Female cocoons were obtained from a commercial breeder (Mauerbienen Shop, Germany). Individuals were visually examined upon emergence to confirm sex. Individual bees were kept in food grade plastic boxes (height 10 cm, radius 6cm). Each bee was provided with ground honeybee collected pollen (General Food Merchants LTD) *ad libitum* as a source of protein and a single 5 ml syringe with either exposure or control feeding solution. Each bee was provided with an unbleached paper ball to hide (∼ 1.5cm radius). Pollen and feeding solution were changed every other day. In addition to clothianidin, sulfoxaflor, and the control, we exposed this species to acetamiprid under the same conditions.

### Lucilia sericata

Fly cocoons were obtained from a commercial breeder (Koppert, UK). Females were identified upon emergence from cocoons by the presence of a larger gap between their eyes in females, compared to the males. After sampling, sex was confirmed by checking for the presence of ovaries. Individual flies were kept in plastic boxes (height 10 cm, radius 6cm). Each fly was provided with pure sucrose ad libitum and a yellow cellulose sponge soaked with either exposure or control feeding solution. Sucrose and exposure solution were changed every other day.

### Vanessa cardui

Butterfly chrysalides were obtained from a commercial breeder (Worldwide Butterflies, UK). Females were recognised in the chrysalis stage by the presence of a distinctive indented line running from the middle of the anal segment to the second segment. After sampling, sex was confirmed by checking for the presence of ovaries. Five butterflies assigned to the same treatment group were marked using water-based markers (POSCA, UK) and kept together in mesh cages (40 cm long, 40 cm wide, 30 cm deep). We assigned the control treatment or one of the two insecticide treatments to all cages in a randomised fashion. Since adult butterflies require artificial feeder training, individuals were hand-fed for 1 minute every day using artificial flowers made from 1.5 ml Eppendorf tubes and yellow or purple vinyl petals. Artificial flowers of both colours and feeding solution-soaked cellulose sponges on petri dishes were always present in the exposure cages and changed every other day. We observed butterflies in all treatments and cages to feed independently after the adjustment phase.

### Bombus terrestris

We acquired ten source colonies of *Bombus terrestris audax* from a commercial breeder (Agralan Growers UK). We transferred the queen, existing brood and 20 workers to wooden boxes (30cm long, 20cm wide, 15cm deep) separated into two equal-sized chambers (foraging and nesting area). We provided each colony with *ad libitum* 30% sucrose solution and organic honeybee-collected pollen (General Food Merchants LTD). We marked all twenty workers using water-based markers (POSCA, UK) and screened the colonies daily for the emergence of new workers. All source colonies were two weeks old when we started setting up the microcolonies. Six callow workers that emerged within 24h were used to assemble microcolonies. We obtained three microcolonies from each source colony.

Each microcolony was kept in a single-chamber wooden box (12cm long, 12cm wide, 10cm deep) and provided with organic pollen *ad libitum* and a single dose of nest substrate (5 parts organic pollen to 1 part 30% sucrose solution in a 2.5 cm Petri dish). We assigned the control treatment or one of the two insecticide treatments to all microcolonies in a randomised fashion. Each microcolony received two 5 mL syringes with either exposure or control feeding solution. We changed pollen and sugar solution every other day.

### Insecticide treatment preparation

Insects were exposed to two insecticides that target nicotinic acetylcholine receptors (nAChRs): clothianidin or sulfoxaflor (Sigma Aldrich, UK). In addition, red mason bees were exposed to acetamiprid (Sigma Aldrich, UK). The use of clothianidin is restricted in the European Union (EFSA 2018), but it is still commonly used worldwide (Simon-Delso et al., 2015). Sulfoxaflor’s use has been increasing (Brown et al., 2016; Siviter et al., 2020) despite recent restrictions within the EU and reports of its high toxicity to bees when applied during flowering (EFSA, 2020). We used the same concentrations for each compound to allow for direct comparisons. We established the concentration at 4.4 ppb which is at the lower end of the concentrations found in pollen and nectar (Botías et al., 2015, 2015; David et al., 2016; Pohorecka et al., 2012; Rundlöf et al., 2015; Siviter et al., 2018) because we aimed to test whether exposure to small concentrations can be detected in the transcriptomic profiles. We prepared insecticide stock solutions by dissolving insecticides in acetone to a concentration of 0.5 mg/mL and stored in darkness at -20℃. Subsequently, we used the stock solution to prepare treatments for each species.

We fed *B. terrestris, O. bicornis,* and *V. cardui* using 30% sucrose solutions as control treatments. For the exposure treatments, we diluted the stock solutions using 30% sucrose solution to 4.4 ppb feeding solutions. Since individuals of *L. sericata* were not attracted to the feeding solution containing only sugar and because young females require protein-rich diet, we prepared amino-acid-rich solutions to feed this species. We prepared 30% solution using sucrose, essential and non-essential amino acids, and cholesterol (Supplementary Table 9), and added the stock solutions to obtain the 4.4 ppb feeding solutions. To avoid insecticide degradation, we stored feeding solutions in darkness at 4℃.

The feeding solutions were made available to the insects *ad libitum* throughout the experiment. Due to biological differences such as body size, metabolic rates, feeding behavior, and nutrient requirements, each species consumed varying amounts of the feeding solutions. This approach of allowing insects to feed ad libitum, rather than providing a standardized amount per individual across species, reflects how species naturally consume resources in their environments. By mirroring their natural feeding behaviours, this method provides more realistic and ecologically valid results, rather than imposing artificial constraints on how much we think they should consume or be exposed to. Notably, no preferential feeding was observed among individuals in different exposure groups.

### Brain tissue dissection, sample processing and sequencing

Bumble bee brains were dissected on dry ice due to their bigger size. In the absence of a queen, one bumble bee worker can become dominant, which induces development of bigger ovaries. We examined abdomens to check for ovarian development and excluded dominant workers from further steps as significant ovarian development may impact gene expression patterns. We randomly selected three non-dominant workers per microcolony and pooled dissected brains for further steps to represent the microcolony level exposure in this social species. We did not pool all six workers per microcolony dues to high mortality rates under exposure to clothianidin. All other insects were dissected in RNAlater solution. Dissected brains were placed in homogenisation tubes with ceramic beads and 400 µl of TRIsol. Samples were placed in -80℃ until homogenisation and RNA extraction. We homogenized the dissected brains in TRIzol using FastPrep96 (45 seconds at 1800 RPM for bumble bees; 35 seconds at 1800 RPM for solitary bees and butterflies; 20 seconds at 1800 RPM for flies because of the variation in tissue size and structure). We isolated RNA using chloroform and purified it with Genaxxon RNA Mini Spin Kit, applying DNase I on-column digestion. We prepared cDNA libraries using the NEBNext® Ultra™ II Directional RNA Library Prep Kit for Illumina. We altered the standard protocol using one-third volumes of enzymes and buffers and 20-200 ng of total RNA input depending on the species with 14 PCR cycles. The library size was quantified using TapeStation 2200 (Agilent, UK) and Qubit 2.0 fluorometer. Libraries were sequenced on NextSeq 500. We obtained the following ranges of 40 bp paired-end reads per sample per species: Bumble bees mean of 36 million (from 22.8 million to 52.2 million reads), solitary bees mean of 22.6 million (from 15.5 million to 30.8 million reads), butterflies mean of 24.3 million (from 17.9 million to 32.8 million reads), flies mean of 32.9 million (from 24.8 million to 39.3 million reads).

### Differential gene expression analysis

We assessed the quality of raw reads using FastQC v0.11.9 (Andrews, 2019). To evaluate alignment qualities, we aligned RNA-seq samples to the reference genomes using STAR v2.7 (Dobin et al., 2013). We used the following reference genomes: Bter_1.0 (PRJNA45869), iOsmBic2.1 (PRJEB44455), ilVanCard2.1 (PRJEB42869), ASM1558622v1 (PRJNA602506). We processed STAR alignments using the RNA-seq module of Qualimap v2.2.1 (Okonechnikov et al., 2016). We summarised the output bam files from STAR using *featureCounts* function in Rsubread v1.22.2 package in R. Each species and exposure group were analysed separately against its corresponding control to enhance statistical power by reducing the complexity of the statistical design relative to the number of replicates per group. This approach also minimizes the risk of batch effects and avoids confounding biological differences that could arise from analysing all species together in a single model. We filtered out genes with relatively low expression levels for each dataset separately adopting the cut-off of a minimum of 10 transcripts in half of the replicates in each of the treatment group. To detect differentially expressed genes between exposure treatments and the control, we applied a Wald test on median-of-ratios normalised counts in DESeq2 v1.32.0 (Love et al., 2014). We included treatment as a factor in all the model designs. For the bumble bees we also included the source colony as a factor. We report genes as differentially expressed using a significance cut-off of 0.05 after false discovery rate adjustment of the Wald test p-values (FDR).

### Correlation of gene expression changes between treatments

We built regression models to evaluate if the directions and intensities of expression changes in the affected genes are comparable between the treatments: 1. Genes recognised as differentially expressed in both treatments; 2. Genes recognised as differentially expressed only under the stronger treatment; 3. A random set of genes not recognised as differentially expressed under any of the treatments. We used the same number of genes in the random set of genes as the number of the overlapping differentially expressed genes between treatments. The rationale was to check whether the direction of the gene expression change, and the intensity were similar between the two treatments. We used log2Fold change values estimated by DESeq2 for all the genes used in the analysis. We applied Pearson correlations and a generalised linear model to assess the correlation strength, slope, and statistical significance of the correlations (Supplementary Table 3).

### Weighted Gene Co-expression Network Analysis (WGCNA)

After obtaining strong correlation values for the overlapping gene sets values in both species, we wanted to test if these genes may belong to the same functional gene groups. We applied weighted gene co-expression network analysis to recognise modules with genes that have similar expression levels. We constructed weighted gene co-expression networks separately for all species using all available samples. We further wanted to detect modules that experience expression shifts under both treatments within each species. We did not detect any modules significantly associated with exposure to sulfoxaflor (the weaker treatment) in the bee species, further showing that sulfoxaflor does not cause major disruptions to the brain transcriptome in bees. We proceeded only with the flies and the butterflies.

To build the gene co-expression network, we first applied the VST as implemented in the DESeq2 package on previously pre-filtered DESeq2 objects for each species. We used the WGCNA v.1-72 package. We applied a soft-thresholding power of 10 for butterflies, and 11 for flies to the correlation matrix based on the scale-free topology fit index of 0.8 and the minimal corresponding mean connectivity of the network value. After constructing the network on all genes, we tested for correlation with exposure treatment: exposed vs. control for each treatment separately using eigengenes and applying Pearson correlation and Benjamini-Hochberg p-value adjustment for multiple testing. We consider modules to be significantly associated with the exposure treatment applying a cut-off of adjusted p-value of 0.05. For the butterflies we detected two modules associated both with exposure to clothianidin and sulfoxaflor, and one more module associated exclusively with sulfoxaflor (the stronger treatment). For the flies, we also detected two modules associated both with exposure to clothianidin and sulfoxaflor, and two more modules associated exclusively with sulfoxaflor. We checked if the genes in modules recognised as associated with both clothianidin and sulfoxaflor exposure overlapped with differentially expressed genes identified by DESeq2. We found higher than expected by random chance overlaps of differentially expressed genes and genes in all the associated modules.

### Orthology search

Using the predicted genes from the reference genomes, we assessed completeness of the longest-isoform protein sequences for each of the four species with BUSCO using the Endopterygota lineage dataset (odb10). The motivation was to demonstrate that, despite the deep divergence times across the species, genome completeness at the Endopterygota lineage levels was sufficient to allow identification of one-to-one orthologs (Supplementary Table 5). Using orthofinder v.2.5.5 and Diamond v.2.1.8 we searched for one-to-one orthologs between all species tested and between pairs of species (Supplementary Table 6).

### KEGG rank-based enrichment analysis

We tested KEGG pathway enrichment using information from KEGG database and the clusterProfiler library in R. We performed this analysis separately for each species and treatment. We excluded both bumble bees and solitary bees exposed to sulfoxaflor because of the low general pattern of differential expression caused by this treatment in both species and close-to-one FDR values for most of the genes. We sorted all genes by FDR values and performed a rank-based test applying a custom function based on the *enrichKEGG* function from clusterProfiles. We applied a cut-off of 0.05 p-value to select pathways with multiple genes affected after exposure and compared the detected pathways between species.

### Comparison to other bumble bee tissues

We extended our analysis to include eight hind leg and eight Malpighian tubule samples from bumble bees originating from the same experimental setup (Witwicka et al., 2024; PRJNA1076820). These samples were processed identically to the brain samples, ensuring consistency. Only unexposed samples were included in the analysis. Surrogate Variable Analysis (SVA) was applied to the VST-normalised data to identify hidden batch effects. The detected batch effect was subsequently removed using the *removeBatchEffect* function in the *limma* v.35.4.2 package. Principal component analysis (PCA) was then performed on the entire dataset. The first principal component (PC1) accounted for 38.3% of the variance, while the second principal component (PC2) accounted for 25.5%. As anticipated, PCA revealed that brain samples from all species, including bumble bees, clustered together, indicating a conserved neural-specific transcriptomic profile across species. In contrast, bumble bee samples from other tissues (hind legs and Malpighian tubules) formed distinct clusters, highlighting tissue-specific differences. This analysis supports the conclusion that neural transcriptomic profiles are more conserved across species than they are to non-neural tissues within a single species (Supplementary Figure 2). We performed this analysis as a form of control to strengthen the validity of our findings, supporting that the observed differences in reactions to insecticide exposure are genuine and not artifacts of the experimental design or technical biases.

### Survival analysis

We collected data on the survival of individual insects throughout the experiments. We fitted Cox proportional hazards regression models to conduct pairwise comparisons of survival between each treatment and the control utilising *survfit* function from the survival v.3.5-7 R library. We performed the test individually for each treatment and species pair. We found the lowest survival rates in flies exposed to both sulfoxaflor (23% survived, P-value < 0.001) and clothianidin (26% survived, P-value < 0.001). We detected lower survival rates under clothianidin in bumble bees (71% survived, P-value < 0.001) and solitary bees (78% survived, P-value < 0.01) compared to sulfoxaflor (98% and 100% survived respectively, P-values > 0.05). All butterfly individuals survived the exposure to both treatments (Supplementary Figure 3).

Interestingly, we observed similar survival rates in flies under both insecticides despite the differences in transcriptomic responses. The survival analysis revealed that flies exposed to sulfoxaflor experiences lethal changes earlier in the exposure with 43% of flies surviving after 4 days of exposure (half time) compared to 94% of flies exposed to clothianidin. Possibly, sulfoxaflor induces more immediate toxic effects, leading to earlier mortality, whereas clothianidin might have a delayed impact in this species. Alternatively, it is possible that some individuals are particularly vulnerable to sulfoxaflor due to intrinsic biological factors, such as differences in metabolism or detoxification capacity. These findings indicate that the timing and mode of action of these insecticides may differ significantly, even when the overall mortality outcomes appear similar. Further investigation is needed to explore these dynamics and their implications for sublethal effects and long-term population impacts.

### Variation in testing power between insect species

The power to detect differentially expressed genes depends on the agreement between the biological replicates within treatment groups and the similarities in genetic background of individuals between groups. Because we controlled for the genetic background of the bumble bees by applying a microcolony-based experimental design, where replicates in all treatments came from the same ten source colonies and were therefore genetically related across treatments, we wanted to test the impact of such adjustment on the detection power of DESeq2 pipeline. We randomly selected control samples from five source colonies and clothianidin samples from the residual five source colonies.

We iterated this process over 100 different selection combinations and built a distribution of the number of differentially expressed genes detected. Next, we repeated this process, but we randomly selected control and clothianidin samples from the same 5 source colonies. We compared both distributions and tested whether it is more likely to obtain higher number of differentially expressed genes when controlling for the source colony effect. When controlling for the colony effect, the median number of differentially expressed genes was 543 compared to 225 when the colony was not accounted for. The distribution when the colony was accounted for was also less positively skewed (skewness = 1.99) compared to when colony was not accounted for (skewness = 0.53). We applied Kolmogorov-Smirnov test to compare the distributions. We detected high numbers of differentially expressed genes using both strategies, however, we conclude that the two distributions are statistically significantly different (D = 0.21, p value = 0.02, Supplementary Figure 4). Therefore, it may be more likely to detect more differentially expressed genes when controlling for the genetic background of the samples. The lower number of detected genes in butterflies and solitary bees may be driven by limited detection power due to underlying genetic differences between individuals, rather than lower insecticide toxicity levels in these species.

We next wanted to measure the agreement between biological replicates within species. The rationale was to see whether in species where individuals were more dissimilar, the power to detect differentially expressed genes was lower. We built a generalised linear model using control samples for all species. We used standard deviation of genes as a response variable and species as explanatory variables. To address issues related to heteroscedasticity and non-normality, we applied variance stabilising transformation (VST) before calculating the standard deviation of gene expression. The use of VST helped to mitigate the presence of extreme values that were observed when standard deviation was calculated on un-normalised counts, which resulted in extremely positively skewed distributions of the residuals and hindered our ability to fit a generalised linear model. Standard deviation of gene expression values after VST is more consistent across genes, as VST reduces the dependence of variance on the mean expression level. Therefore, if we still see differences in standard deviation between species calculated using VST-normalised counts, the differences in standard deviation between genes in non-normalised counts should be greater. To assure the best model fit for continuous data with a positively skewed distribution, we applied the gamma distribution. The outputs showed the greatest differences between fly samples compared to the other three species (all p-values < 0.001), indicating greater variability between the fly individuals. Such pattern shows that despite differences between individuals, both insecticides impacted the flies so much that we were able to detect high numbers of differentially expressed genes, compared to other species. This strong impact of exposure was also visible in the low survival rates in this species compared to other species used in this study.

Given the above observations, we do not use the number of differentially expressed genes to compare the toxicity levels of insecticides between species, and only provide a quantitative measure of which insecticide affects each species more strongly: While clothianidin caused relatively more changes in both bee species compared to sulfoxaflor, this pattern was reversed in non-bee species, where sulfoxaflor caused more changes in the transcriptome compared to clothianidin.

