## Supplementary Figures for "Minimal overlaps in responses to insecticides between pollinator species"

**Supplementary Table 1.** Number of down- and up-regulated genes detected by DESeq2 analysis (FDR < 0.05). Each species was analysed individually, with two DESeq2 models built per species, comparing each pesticide treatment to the control.

| | | Down-regulated | Up-regulated | Unchanged |
|---|---|---|---|---|
| Sulfoxaflor | Fly | 753 | 1115 | 14568 |
| | Butterfly | 428 | 306 | 14266 |
| | Solitary bee | 3 | 1 | 12626 |
| | Bumble bee | 1 | 0 | 12007 |
| Clothianidin | Fly | 82 | 416 | 15938 |
| | Butterfly | 40 | 12 | 14948 |
| | Solitary bee | 367 | 392 | 11871 |
| | Bumble bee | 1019 | 1167 | 9822 |

**Supplementary Table 2.** Percentage of differentially expressed genes in proportion to the total number of genes in the genome. The scale of transcriptomic changes following exposure varied considerably across species, with flies and bumble bees exhibiting greater transcriptomic changes compared to butterflies and solitary bees. However, this pattern may reflect differences in statistical testing power rather than true biological effects.

| | | % of differentially expressed genes | |
|---|---|---|---|
| | No. genes | Sulfoxaflor | Clothianidin |
| Fly | 16436 | **11.37%** | **3.03%** |
| Butterfly | 15000 | **4.89%** | **0.35%** |
| Solitary bee | 12630 | **0.03%** | **6%** |
| Bumble bee | 12008 | **0%** | **18.20%** |

**Supplementary Table 3.** Models illustrating the relationship between the direction and magnitude of gene expression changes influenced by two distinct insecticides. The table presents the Pearson correlation coefficient (*r*), and the slope derived from generalized linear models (GLM). Comparisons of overlapping differentially expressed genes (DEGs) reveal a highly proportional correlation between expression changes in butterflies and flies, as reflected by slopes close to 1 and strong *r* values. Genes differentially expressed only under sulfoxaflor also show altered expression under clothianidin but to a lesser extent, leading to a less proportional correlation. For solitary bees, most genes differentially expressed under acetamiprid are also differentially expressed under clothianidin, maintaining a proportional relationship. In bumble bees, the gene expression changes induced by imidacloprid and clothianidin are also strongly correlated, emphasizing a consistent relationship across the insecticides. Significance codes: * < 0.05; ** < 0.01; *** < 0.001.

| **Comparison** | | *t-value* | *DFs* | **Pearson *r*** | **GLM *slope*** | *SE* | *p-value* |
|---|---|---|---|---|---|---|---|
| *butterfly* | *overlaping genes [sulfoxaflor & clothianidin]* | 36.21 | 40.00 | **0.99** | **0.98** | 0.03 | *** |
| *fly* | *overlaping genes [sulfoxaflor & clothianidin]* | 64.00 | 423.00 | **0.95** | **0.99** | 0.02 | *** |
| *butterfly* | *sulfoxaflor only DEGs* | 48.17 | 690.00 | **0.88** | **1.51** | 0.03 | *** |
| *fly* | *sulfoxaflor only DEGs* | 87.30 | 1441.00 | **0.92** | **1.31** | 0.02 | *** |
| *solitary bee* | *acetamiprid DEGs* | 25.00 | 26.00 | **0.98** | **1.31** | 0.05 | *** |
| *bumble bee* | *overlaping genes [imidacloprid & clothianidin]* | 19.00 | 146.00 | **0.84** | **0.94** | 0.05 | *** |

**Supplementary Table 4.** WGCNA modules detected in the fly *L. sericata* and the number of genes in each module paired with the number of genes detected as differentially expressed in each module. Two of the modules (X and XIV) were detected as statistically significantly associated with pesticide exposure under both treatments. In the fly, 68% of the differentially expressed genes under sulfoxaflor and 87% under clothianidin exposure were found within these modules. Significance codes: * < 0.05; ** < 0.01; *** < 0.001.

| Module: | No. genes: | No. genes differentially expressed: | |
|---|---|---|---|
| | | Sulfoxaflor | Clothianidin |
| I | 20 | 1 | 1 |
| II | 26 | 3 | 1 |
| III | 33 | 0 | 0 |
| IV | 37 | 0 | 0 |
| V | 53 | 3 | 0 |
| VI | 57 | 0 | 0 |
| VII | 122 | 0 | 0 |
| VIII | 130 | 0 | 2 |
| **IX** | **142** | ***46 **** | 1 |
| **X** | **837** | ***436 ***** | ***48 **** |
| **XI** | **853** | ***357 ***** | 4 |
| XII | 1097 | 2 | 0 |
| XIII | 2268 | 21 | 4 |
| **XIV** | **2649** | ***829 **** | ***386 **** |

**Supplementary Table 5.** WGCNA modules detected in the butterfly *V. cardui* and the number of genes in each module paired with the number of genes detected as differentially expressed in each module. Two of the modules (XIII and XIV) were detected as statistically significantly associated with pesticide exposure under both treatments. In the butterfly, 33% of the differentially expressed genes under sulfoxaflor and 27% under clothianidin exposure were in these modules. Notably, the modules detected in the butterfly were smaller. Significance codes: . = 0.05; * < 0.05; ** < 0.01; *** < 0.001.

| Module: | No. genes: | No. genes differentially expressed: | |
|---|---|---|---|
| | | Sulfoxaflor | Clothianidin |
| I | 21 | 0 | 0 |
| II | 22 | 0 | 0 |
| III | 22 | 0 | 0 |
| IV | 23 | 0 | 0 |
| V | 26 | 0 | 0 |
| VI | 27 | 0 | 0 |
| VII | 27 | 0 | 0 |
| VIII | 27 | 0 | 0 |
| IX | 27 | 0 | 0 |
| X | 33 | 0 | 0 |
| XI | 42 | 8 | 1 |
| XII | 130 | 11 | 0 |
| **XIII** | **199** | ***136 ****** | ***9 **** |
| **XIV** | **285** | ***107 ****** | ***6 .*** |
| XV | 357 | 3 | 0 |
| XVI | 507 | 16 | 1 |
| **XVII** | **563** | ***171 ***** | 2 |
| XVIII | 2001 | 66 | 5 |

**Supplementary Table 6.** The BUSCO genome completeness scores for the Endopterygota lineage.

| | Bumble bee | Solitary bee | Butterfly | Fly |
|---|---|---|---|---|
| Complete BUSCOs: | 2111 (99.4%) | 2104 (99.1%) | 2115 (99.6%) | 2061 (97%) |
| Complete and single-copy BUSCOs: | 1710 (80.5%) | 1659 (78.1%) | 1960 (92.3%) | 1682 (79.2%) |
| Complete and duplicated BUSCOs: | 401 (18.9%) | 445 (21%) | 155 (7.3%) | 379 (17.8%) |
| Fragmented BUSCOs: | 3 (0.1%) | 7 (0.3%) | 4 (0.2%) | 16 (0.8%) |
| Missing BUSCOs: | 10 (0.5%) | 13 (0.6%) | 5 (0.2%) | 47 (2.2%) |

**Supplementary Table 7.** Number of one-to-one orthologs detected between the four species.

| | Fly | Butterfly | Solitary bee | Bumble bee |
|---|---|---|---|---|
| Fly | 16436 | | | |
| Butterfly | 4816 | 15000 | | |
| Solitary bee | 4103 | 5181 | 12630 | |
| Bumble bee | 4221 | 5287 | 6141 | 12008 |
| All species | 2965 | | | |

**Supplementary Table 8.** Overlaps of differentially expressed one-to-one orthologs between species. Purple text indicates exposure to clothianidin, yellow text indicates exposure to sulfoxaflor. The only overlap that was statistically significant was between the two bee species. However, the direction (up- or down-regulated) of expression of the overlapping differentially expressed one-to-one orthologs varied between the two species with only 25 genes exhibiting similar expression patterns. Expected minimum overlap is drown from hypergeometric distributions based on the number of one-to-one orthologs between a species pair and the number of differentially expressed one-to-one orthologs in these species. In all comparisons the observed values were lower than the minimal expected overlaps.

| | | *No. differentially expressed genes* | | | *No. differentially expressed one-to-one orthologs* | | *Percentage* | | | | | |
|---|---|---|---|---|---|---|---|---|---|---|---|---|
| *Species 1* | *Species 2* | *Species 1* | *Species 2* | *No. one-to-one orthologs* | *Species 1* | *Species 2* | *Species 1* | *Species 2* | *Overlap* | *Overlap of genes with the same direction of change* | | *Expected minimum overlap* |
| Solitary bee | Bumble bee | 759 | 2186 | 6141 | **454** | **1244** | 59.82 | 56.91 | 136 | **25** | < | **93** |
| Butterfly | Bumble bee | 734 | 2186 | 5287 | **360** | **1129** | 49.05 | 51.65 | 75 | **23** | < | **76** |
| Solitary bee | Butterfly | 759 | 734 | 5181 | **387** | **350** | 50.99 | 47.68 | 23 | **8** | < | **27** |
| Fly | Butterfly | 1868 | 734 | 4816 | **667** | **327** | 35.71 | 44.55 | 54 | **25** | < | **45** |
| Fly | Bumble bee | 1868 | 2186 | 4221 | **599** | **925** | 32.07 | 42.31 | 151 | **108** | < | **130** |
| Solitary bee | Fly | 759 | 1868 | 4103 | **290** | **603** | 38.21 | 32.28 | 42 | **21** | < | **44** |
| Solitary bee | Butterfly | 759 | 52 | 5181 | **387** | **22** | 50.99 | 42.31 | 3 | **1** | < | **3.5** |
| Butterfly | Bumble bee | 52 | 2186 | 5287 | **30** | **1129** | 57.69 | 51.65 | 5 | **1** | < | **8.5** |
| Solitary bee | Fly | 759 | 498 | 4103 | **290** | **77** | 38.21 | 15.46 | 15 | **8** | < | **11** |
| Fly | Bumble bee | 498 | 2186 | 4221 | **73** | **925** | 14.66 | 42.31 | 13 | **7** | < | **18** |
| Fly | Butterfly | 498 | 52 | 4816 | **86** | **25** | 17.27 | 48.08 | 0 | **0** | < | **3** |

**Supplementary Table 9.** *Lucillia sericata* food composition. We prepared a 30% solution by mixing glucose, essential and non-essential amino acids, cholesterol, and water. This solution served as the control feeding solution and was also used to dilute the pesticide stock solution to 4.4 ppb.

| *Lucilia sericata:* diet proportions | | [g or mL] |
|---|---|---|
| Glucose | | 24 |
| EAA | Arginine | 0.3 |
| | Histidine | 0.3 |
| | Isoleucine | 0.3 |
| | Leucine | 0.3 |
| | Lysine | 0.3 |
| | Methionine | 0.3 |
| | Phenylalanine | 0.3 |
| | Threonine | 0.3 |
| | Tryptophan | 0.3 |
| | Valine | 0.3 |
| N-EAA | Tyrosine | 0.3 |
| | Alanine | 0.3 |
| | Asparagine | 0.3 |
| | Aspartic acid | 0.2 |
| | Cysteine | 0.3 |
| | Glutamic acid | 0.3 |
| | Glutamine | 0.3 |
| | Glycine | 0.3 |
| | Proline | 0.3 |
| | Serine | 0.3 |
| Cholesterol | | 0.1 |
| Water | | 70 |

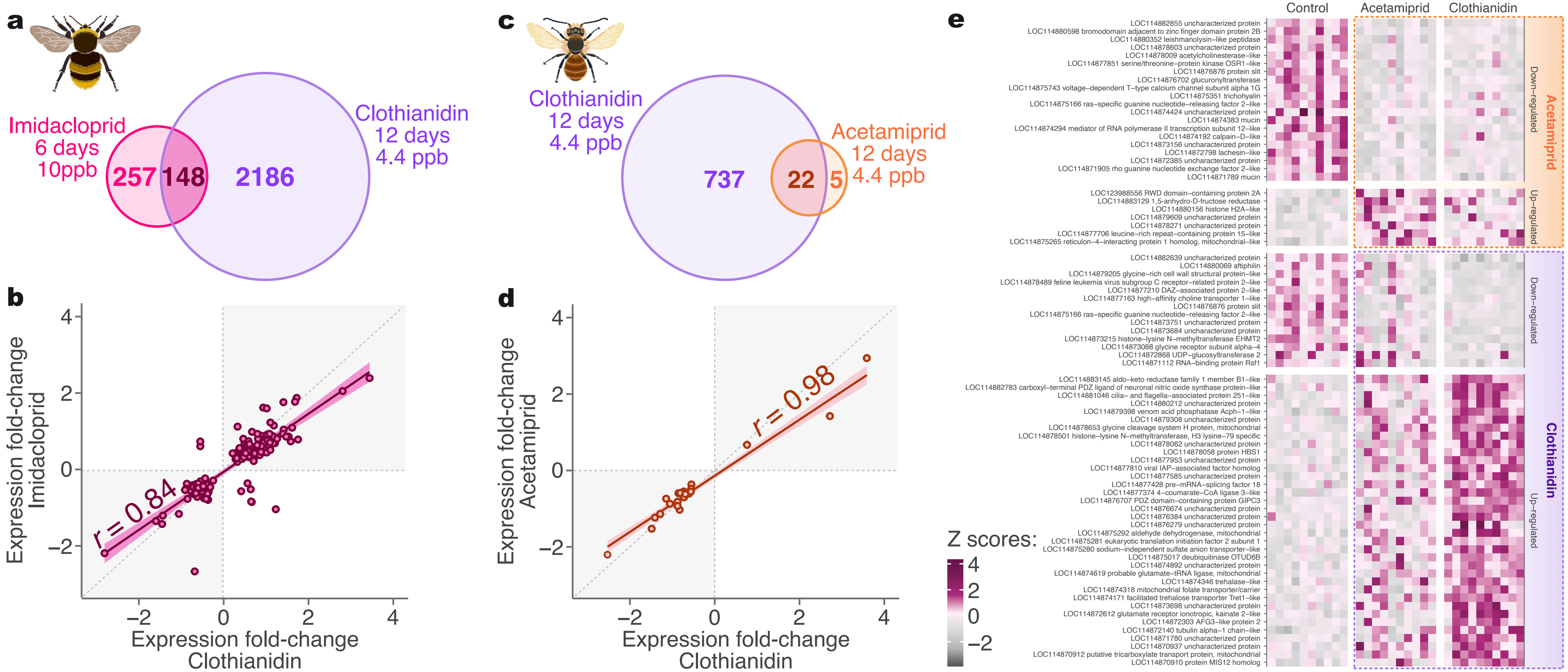


**Supplementary Figure 1.** Since sulfoxaflor did not induce notable transcriptomic changes in the bee species, we conducted additional comparisons with other insecticides. **A.** We compared the differentially expressed genes detected under exposure to imidacloprid and clothianidin in bumble bees. The imidacloprid gene list was derived from Bebane et al. (2019), where bees were exposed to 10 ppb for six days, compared to the 4.4 ppb over 12 days used in this study. Despite differences in exposure conditions, we observed an overlap of 148 differentially expressed genes between the two compounds. The dose and time of exposure have greater effect of the exposure outcome than the insecticide type itself (Witwicka et al., 2024), additionally, differences in experimental design and data processing likely reduce the strength of the correlation compared to other species. Despite this, a statistically significant overlap was still observed (hypergeometric test P-value < 0.001). **B.** The amplitudes of expression changes in the overlapping differentially expressed genes were strongly correlated between imidacloprid and clothianidin (*r* = 0.84). **C.** We exposed solitary bees to acetamiprid and compared their DEGs with those detected under clothianidin exposure. We identified 27 differentially expressed genes in response to acetamiprid, 22 of which overlapped with differentially expressed genes detected under clothianidin exposure (hypergeometric test P-value < 0.001). **D.** The expression amplitudes of the overlapping genes were highly correlated between acetamiprid and clothianidin exposure (*r* = 0.98), indicating similar transcriptomic effects across these compounds. **E.** Heatmap showing all differentially expressed genes detected under acetamiprid exposure and the top 50 genes with the lowest FDR values under clothianidin exposure. Gene expression patterns in bees exposed to acetamiprid and clothianidin were more similar to each other than to control samples, underscoring the shared molecular mechanisms between these insecticides.

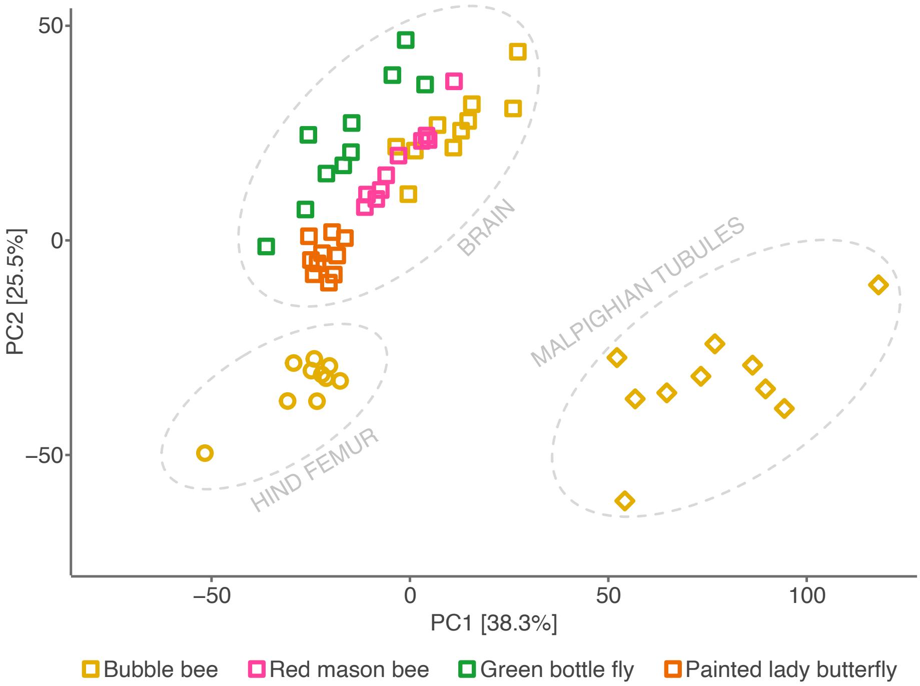


**Supplementary Figure 2.** PCA showing the clustering of unexposed samples by tissue based on one-to-one orthologs present in all species. Brain samples from different species cluster more closely compared to two other tissues from the bumble bee. This clustering highlights the conserved molecular profile of brain tissue across species. Bumble bee tissue data from Witwicka et al., 2024 (PRJNA1076820).

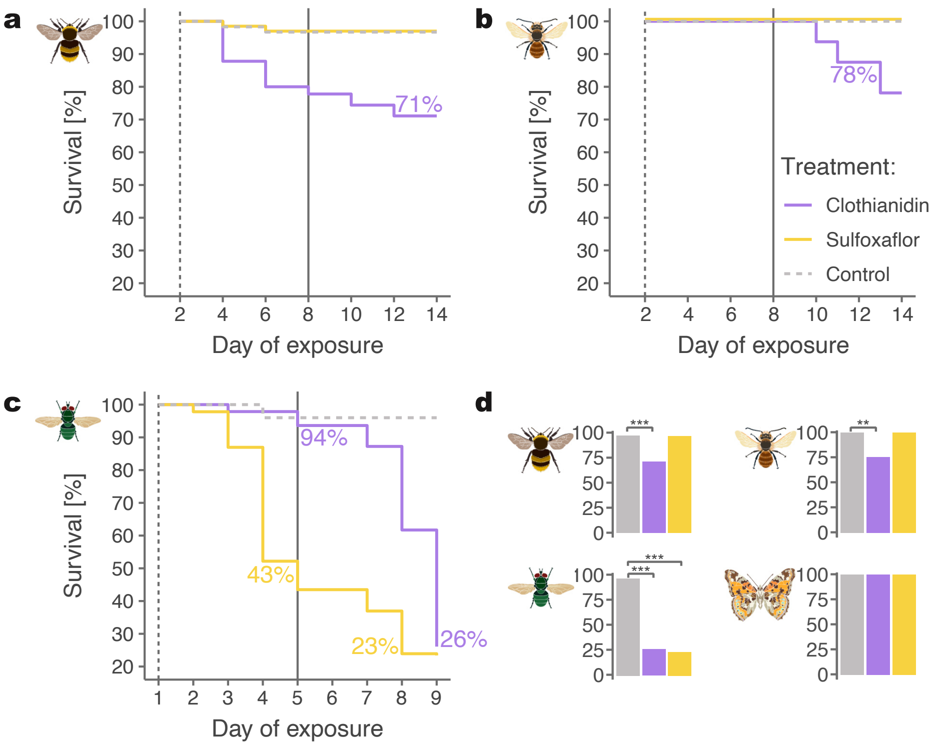


**Supplementary Figure 3.** Kaplan-Meier plots showing the survival rates of **A.** bumble bees, **B.** red mason bees, **C.** green bottle flies under exposure to clothianidin and sulfoxaflor. In bees sulfoxaflor caused no mortality compared to the control. In flies sulfoxaflor and clothianidin caused comparable mortality rates at the end of the experiments while sulfoxaflor (the stronger compound) caused more immediate lethal effects with 43% surviving to the middle of exposure compared to 23% under clothianidin exposure. **D.** Total mortality rates across species at the end of exposure. All butterflies exposed to clothianidin and sulfoxaflor survived the exposure. Significance codes for the Cox proportional hazards regression models: * < 0.05; ** < 0.01; *** < 0.001.

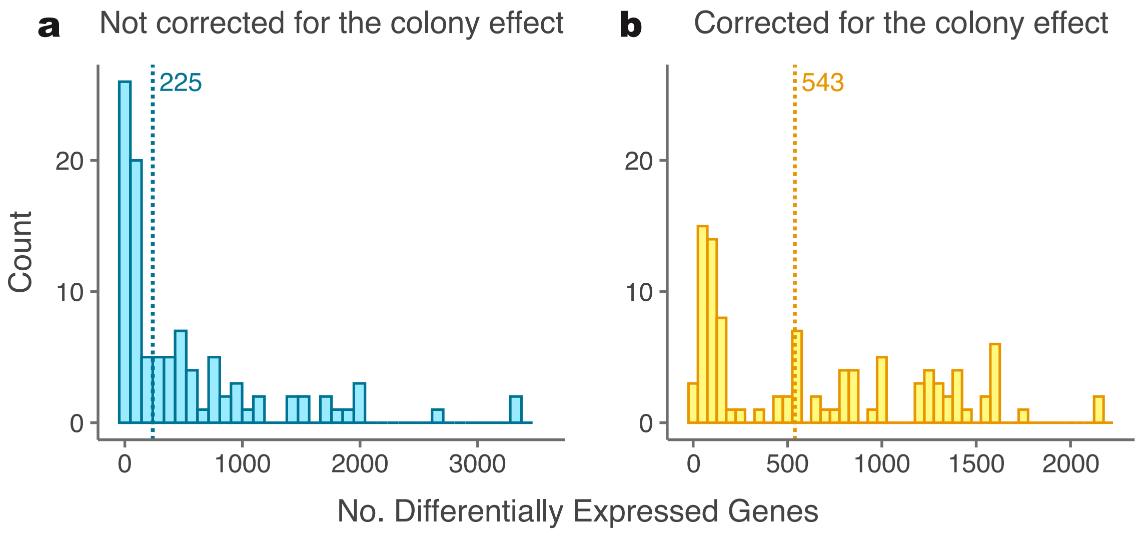


**Supplementary Figure 4.** Number of differentially expressed genes **A.** between bumble bee control and clothianidin-exposed samples corrected for the colony effect and **B.** with no correction for colony effect. Using bumble bees from the same source colonies increases likelihood of differential gene detection.

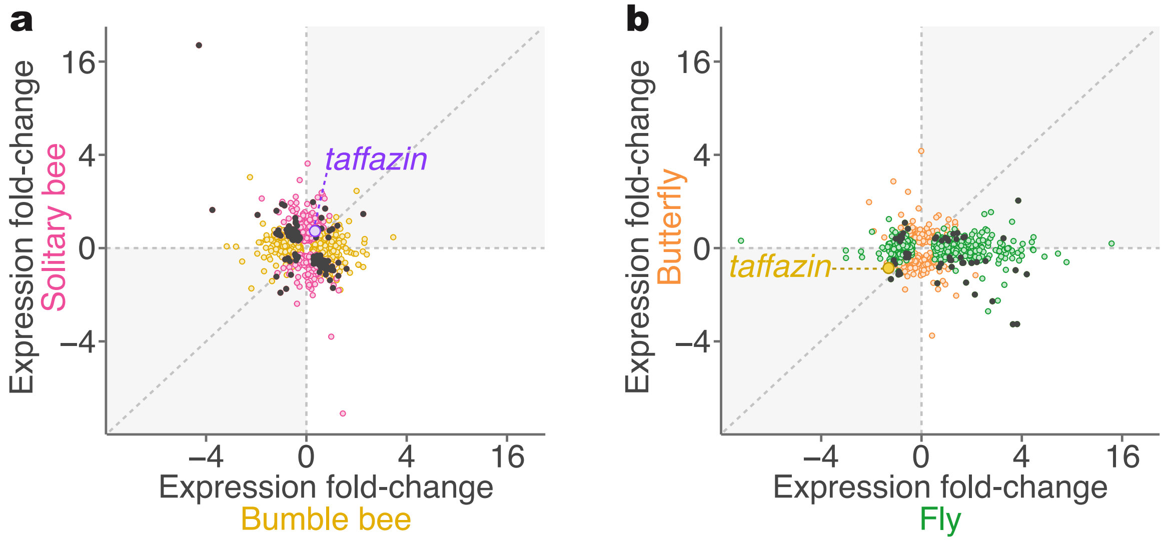


**Supplementary Figure 5.** Correlation of the magnitude of expression fold-change in one-to-one orthologs in **A.** bumble bees and red mason bees exposed to clothianidin, and **B. in** flies and butterflies exposed to sulfoxaflor. This figure mirrors the plots in Figure 3 but includes outlier genes, which were removed in the main figure for clarity.
